# Loss-of-function tolerance of enhancers in the human genome

**DOI:** 10.1101/608257

**Authors:** Duo Xu, Omer Gokcumen, Ekta Khurana

## Abstract

Previous studies have surveyed the potential impact of loss-of-function (LoF) variants and identified LoF-tolerant protein-coding genes. However, the tolerance of human genomes to losing enhancers has not yet been evaluated. Here we present the catalog of LoF-tolerant enhancers using structural variants from whole-genome sequences. Using a conservative approach, we estimate that each individual human genome possesses at least 28 LoF-tolerant enhancers on average. We assessed the properties of LoF-tolerant enhancers in a unified regulatory network constructed by integrating tissue-specific enhancers and gene-gene interactions. We find that LoF-tolerant enhancers are more tissue-specific and regulate fewer and more dispensable genes. They are enriched in immune-related cells while LoF-intolerant enhancers are enriched in kidney and brain/neuronal stem cells. We developed a supervised learning approach to predict the LoF-tolerance of enhancers, which achieved an AUROC of 96%. We predict 5,677 more enhancers would be likely tolerant to LoF and 75 enhancers that would be highly LoF-intolerant. Our predictions are supported by known set of disease enhancers and novel deletions from PacBio sequencing. The LoF-tolerance scores provided here will serve as an important reference for disease studies.

## Introduction

Loss-of-function (LoF) variants in genes are defined as those which impair or eliminate the function of the encoded protein. Despite their protein-coding disruption, it has been shown that some LoF variants can be tolerated in healthy individuals (Ng et al. 2008, Pelak et al. 2010, Genomes Project et al. 2010, Telenti et al. 2016). Genes harboring homozygous LoF variants are called LoF-tolerant genes. Multiple studies have shown the average number of LoF variants ranges from 100∼200 per individual (MacArthur et al. 2012, Genomes Project et al. 2015, Lek et al. 2016). In addition, MacArthur et al estimated that on average there are 20 LoF-tolerant genes per human genome (MacArthur et al. 2012). Such lists of LoF variants have greatly aided gene prioritization in disease studies by providing functional references for variants (Kathiresan and Srivastava 2012, Tg and Hdl Working Group of the Exome Sequencing Project et al. 2014, Gilissen et al. 2014, Genovese et al. 2016, Yu et al. 2018). It also enabled estimations of gene indispensability by providing a confident set of LoF variants and LoF-tolerant genes in human genomes (Khurana et al. 2013, MacArthur et al. 2012).

However, in stark contrast to protein-coding genes, our knowledge about the dispensability of non-coding regulatory elements is limited. The atlas of cell- and tissue-specific regulatory elements developed by large-scale efforts, such as ENCODE (Consortium 2012, Davis et al. 2018), Roadmap Epigenomics Mapping Consortium (Roadmap Epigenomics et al. 2015), FANTOM (Andersson et al. 2014) and the availability of thousands of whole-genomes makes this an opportune time to ask the same questions that were asked for protein-coding genes and to identify the non-coding elements that can tolerate homozygous LoF.

It has been shown that ‘shadow’ enhancers, defined as the ones that have similar functions to the proximal primary enhancers but locate at distal locations, can be deleted in Drosophila without affecting the viability (Perry et al. 2010). A recent study showed that deletion of some individual enhancers did not significantly affect the fitness of mice, but deletion of pairs of enhancers regulating the same gene led to abnormal limb development (Osterwalder et al. 2018). Thus, the phenotypic effect stemming from the loss of a single enhancer may be mitigated by the activity of another enhancer, whose function is redundant to the deleted one, and is therefore only apparent if both enhancers are deleted. This apparent redundancy of enhancers is hypothesized to provide robustness in gene expression in response to the fluctuating environmental conditions (Macneil and Walhout 2011, Wunderlich et al. 2015). These studies suggest that enhancers can act redundantly in groups instead of stand-alone units. On the other hand, alterations at single enhancers have been implicated in rare Mendelian diseases (Ghiasvand et al. 2011, Albuisson et al. 2011, Weedon et al. 2014, Kapoor et al. 2014, Kremer et al. 2017) and genome-wide association studies (GWAS) have found that many susceptibility loci for common diseases reside in enhancers (MacArthur et al. 2017, Trynka et al. 2013, Hindorff et al. 2009, Maurano et al. 2012, Wang et al. 2018). It is expected that loss of essential enhancers would have strong fitness consequences, while LoF-tolerant enhancers would lie at the other end of the spectrum and their loss would not elicit substantial phenotypic impact. Thus, it is important to have a prioritization scheme for LoF-tolerance vs. disease-causing potential of enhancers based on their essentiality. However, due to the redundancy and complexity of tissue-specific regulatory networks, such prioritization of enhancers has long remained a challenging task.

Here we report a systematic computational approach that uses machine learning to predict the LoF-tolerance of all enhancers in the human genome. We built an integrated regulatory network, MegaNet, in which the nodes consist of enhancers and genes. The edges between enhancers and genes correspond to tissue-specific regulation and those between genes include protein-protein (Stark et al. 2006), metabolic (Kanehisa et al. 2010), phosphorylation (Lin et al. 2010) and signaling interactions (Korcsmáros et al. 2010). We used deletions from 2,054 whole-genomes to identify the LoF-tolerant enhancers in this network while taking ultra-conserved enhancers with experimentally validated enhancer activity as LoF-intolerant (Bejerano et al. 2004, Dickel et al. 2018, Visel et al. 2007, Visel et al. 2008). We used the characteristic differences between LoF-tolerant and LoF-intolerant enhancers in MegaNet to build a random forest model to predict the LoF-tolerance of all enhancers in the human genome. The LoF-tolerance scores of enhancers provided in this study can significantly facilitate the interpretation and prioritization of non-coding sequence variants for disease and functional studies.

## Results

### Construction of MegaNet

Integration of transcription factor (TF) binding profiles, chromatin features and expression data has revealed the architecture of regulatory networks (He et al. 2014, Yip et al. 2012, Zhu et al. 2016, Whalen, Truty and Pollard 2016, Roy et al. 2016). Availability of tissue-specific annotations has also enabled the construction of tissue-specific regulatory networks (Cao et al. 2017). In order to systematically evaluate the LoF-tolerance of enhancers in tissue-specific regulatory networks, we collected 246,028 unique enhancers regulating 19,170 genes from enhancer-target networks (Cao et al. 2017). We constructed an integrated mega network (MegaNet) for joint assessment of the enhancer properties in the enhancer-gene regulation networks (Cao et al. 2017) and gene centrality in the gene-gene interaction networks (Khurana et al. 2013). The gene-gene interactions in MegaNet consist of protein-protein (Stark et al. 2006), metabolic (Kanehisa et al. 2010), phosphorylation (Lin et al. 2010) and signaling interactions (Korcsmáros et al. 2010).

In the MegaNet, enhancers and genes represent the two kinds of nodes. The directed regulation from enhancers to genes and the undirected interactions between genes are the edges. In order to annotate the tissue-specific properties of nodes and edges in the MegaNet, the enhancer->gene regulation edges are weighted by the number of tissues in which they are active and annotated by tissue types (Figure 1a).

**Figure 1.**
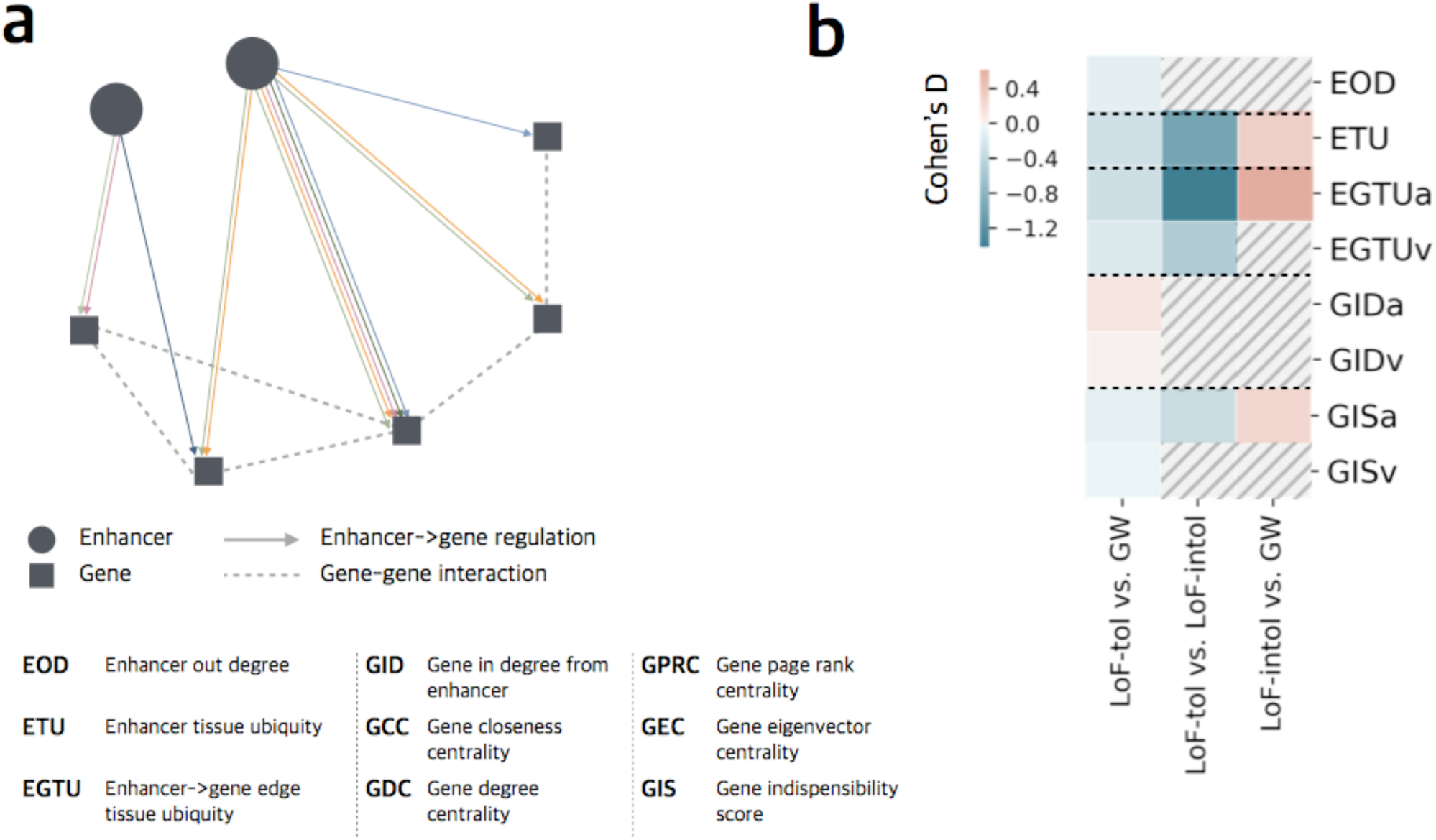
MegaNet features. a) Schema of the MegaNet, circle and square represent nodes for enhancers and genes, respectively, and colored directed arrows are enhancer->gene regulation edges. Different colors represent the interactions active in different tissues. Dashed lines represent the gene-gene interactions. b) Only significant comparisons (P-value < 0.05) are shown in color, non-significant ones are marked by dashed lines. Color scale represents Cohen’s D for effective size, positive values stand for higher average while negative values stand for lower average. LoF-tol, LoF-intol and GW represent LoF-tolerant, LoF-intolerant and genome-wide respectively. ‘a’ and ‘v’ stand for the average and variance for the corresponding features in a).

### LoF-tolerant enhancers

We adopted the enhancers annotated by Cao et al. (Cao et al. 2017, Methods) which were collected from the ENCODE and Roadmap Epigenomics projects (Consortium 2012, Roadmap Epigenomics et al. 2015). Since samples in the 1000 Genomes Project consist of individuals without strong disease phenotypes (Genomes Project et al. 2010, Auton et al. 2015), we define enhancers that can be homozygously deleted in those individuals as LoF-tolerant enhancers. This is similar to the approach used previously for identification of LoF-tolerant genes (Ng et al. 2008, Pelak et al. 2010, MacArthur et al. 2012). More specifically, to identify the LoF-tolerant enhancers, we identified deletions which occur homozygously in at least one individual among the 2,504 from the 1000 Genomes Project (Sudmant et al. 2015) and intersected them with enhancers. In order to avoid bias introduced by protein-coding regions, deletions that overlap coding exons were excluded. While deletion of parts of enhancers may also lead to loss of their activity, we used a conservative estimation of LoF-tolerant enhancers by only including those that are completely deleted in a homozygous manner. In line with this, our approach also does not include LoF of enhancers by SNVs due to the difficulties in predicting their functional impact. In total, 886 enhancers are identified as LoF-tolerant. The number of LoF-tolerant enhancers per individual genome ranges from 8 to 78 (Supplementary Figure 1).

### LoF-intolerant enhancers

Efforts to identify enhancers that are intolerant to LoF have relied on evolutionary conservation to identify the ultra-conserved non-coding elements in the genome (Bejerano et al. 2004). 256 ultra-conserved non-exonic elements have been identified by absolute conservation between orthologous regions of the human, rat and mouse genomes. While the initial study to probe the indispensability of ultra-conserved enhancers showed that their deletion does not affect the viability of mice (Ahituv et al. 2007), more recent studies have found that the mice suffer from severe developmental defects (Dickel et al. 2018), indicating that ultra-conserved enhancers are in fact LoF-intolerant as their loss strongly adversely affects organismal fitness. Overall, we compiled 49 LoF-intolerant enhancers, which correspond to the ultra-conserved non-coding elements that have shown enhancer activity by consistent reporter gene expression in at least three transgenic mice embryos (Bejerano et al. 2004, Visel et al. 2007, Visel et al. 2008, Dickel et al. 2018). Furthermore, in agreement with previous studies (Katzman et al. 2007), we observe a depletion of common polymorphisms and an enrichment of rare variants at LoF-intolerant enhancer regions, providing additional support for negative selection preventing mutations accumulating in LoF-intolerant enhancers (Supplementary Figure 2).

### Properties of LoF-tolerant and -intolerant enhancers in the MegaNet

We analyzed the properties of enhancers in MegaNet using enhancer out-degree (EOD, number of genes that an enhancer targets), enhancer tissue ubiquity (ETU, total number of tissues the enhancer is active in), and enhancer->gene edge tissue ubiquity (EGTU, the number of tissues in which the edges are active) (detailed feature description provided in Supplementary Table 1). ETU describes the total number of tissues that the enhancer is active in, while EGTU describes the number of tissues that an enhancer->gene regulation edge is active in (Figure 1a). Khurana et al. integrated multiple biological networks to evaluate the functional essentiality of genes in the human genome (Khurana et al. 2013). We assigned the gene indispensability scores generated from that study to genes in our network to integrate the gene indispensability (GIS) in the MegaNet. In order to assess the enhancer-gene interaction landscape in the MegaNet, we also calculated the number of enhancers regulating each gene (Gene In-Degree, GID), and other network centrality metrics as additional gene properties (detailed feature description provided in Supplementary Table 1). Due to the characteristic architecture of regulatory networks, an enhancer can regulate multiple genes and a gene can be regulated by multiple enhancers as well. Enhancers regulating multiple genes will have multiple values for each gene feature. We consider both the mean and variance to represent their values, and they are represented with an extension “a” (average) or “v” (variance). For example, the enhancer on the left in Figure 1a regulates two genes in three different tissues. The ETU of the enhancer is 3 while the EGTU is a collection of (2,1). The EGTUa for the enhancer will be 1.5 and EGTUv will be 0.25 (Methods).

#### LoF-tolerant enhancers tend to be tissue-specific and regulate fewer, more dispensable genes

We compared the network properties of LoF-tolerant and LoF-intolerant enhancers and genome-wide expectation (GW, all other enhancers in the MegaNet). We find that LoF-tolerant enhancers regulate significantly fewer genes (i.e., they have lower EOD) compared to genome-wide expectation and are active in fewer tissues (ETU) compared to both genome-wide expectation and LoF-intolerant enhancers (Figure 1b, Supplementary Figure 3a). In addition, genes regulated by LoF-tolerant enhancers are more dispensable (lower average gene indispensability score, GISa) compared to genome-wide expectation and LoF-intolerant enhancers. In order to account for enhancers with the same average EGTU, but different variance, we also analyzed the variance of EGTU. Both average edge tissue ubiquity (EGTUa) and its variance (EGTUv) are lower for LoF-tolerant enhancers, indicating that their interactions tend to be more tissue-specific (Figure 1b). Overall, these observations indicate that LoF-tolerant enhancers are in general less versatile in the genome and tend to target specific genes in specific tissues.

#### Genes regulated by LoF-tolerant enhancers are regulated by more enhancers

Interestingly, we observe that the genes that LoF-tolerant enhancers regulate, have significantly more enhancers regulating them (higher Average Gene In-degree, GIDa) (Figure 1b). As mentioned in the Introduction, enhancers can act in groups rather than as single units (Perry et al. 2010, Macneil and Walhout 2011, Wunderlich et al. 2015, Osterwalder et al. 2018). Here, we show that this trend exists in a genome-wide manner, such that enhancers targeting genes that are regulated by multiple enhancers tend to be more LoF-tolerant. Thus, LoF-tolerant enhancers potentially function redundantly to prevent severe phenotypic effects when one or more enhancers are lost.

#### LoF-intolerant enhancers are enriched in kidney and brain/neuronal stem cells while LoF-tolerant enhancers are enriched in immune related cells

Furthermore, to analyze the tissue-specific properties of enhancers, we extracted the tissue-specific sub-networks from the MegaNet. We observe that different tissues exhibit differential enrichment of LoF-tolerant vs. LoF-intolerant enhancers. We calculated the odds ratio of LoF-tolerant and -intolerant enhancers for each tissue compared to the total number of LoF-tolerant and -intolerant enhancers across all other tissues respectively (Figure 2). We found that the proportion of LoF-intolerant enhancers in kidney and neuronal stem cell/brain tissues is significantly enriched (Fisher’s exact test P-value = 0.010 and 2.80e-11 respectively, Figure 2). Interestingly, this trend is reversed in cells involved in immune response (Hematopoietic stem cells (HSC) & B-cell and T-cell), where LoF-intolerant enhancers are depleted while LoF-tolerant are enriched (Fisher’s exact test P-value = 4.94e-4 and 1.70e-7, Figure 2). Our results are consistent with the previous knowledge that ultra-conserved enhancers are related to brain or developmental function (Dickel et al. 2018). Importantly, they show that enhancers involved in immune response tend to be more LoF-tolerant.

**Figure 2.**
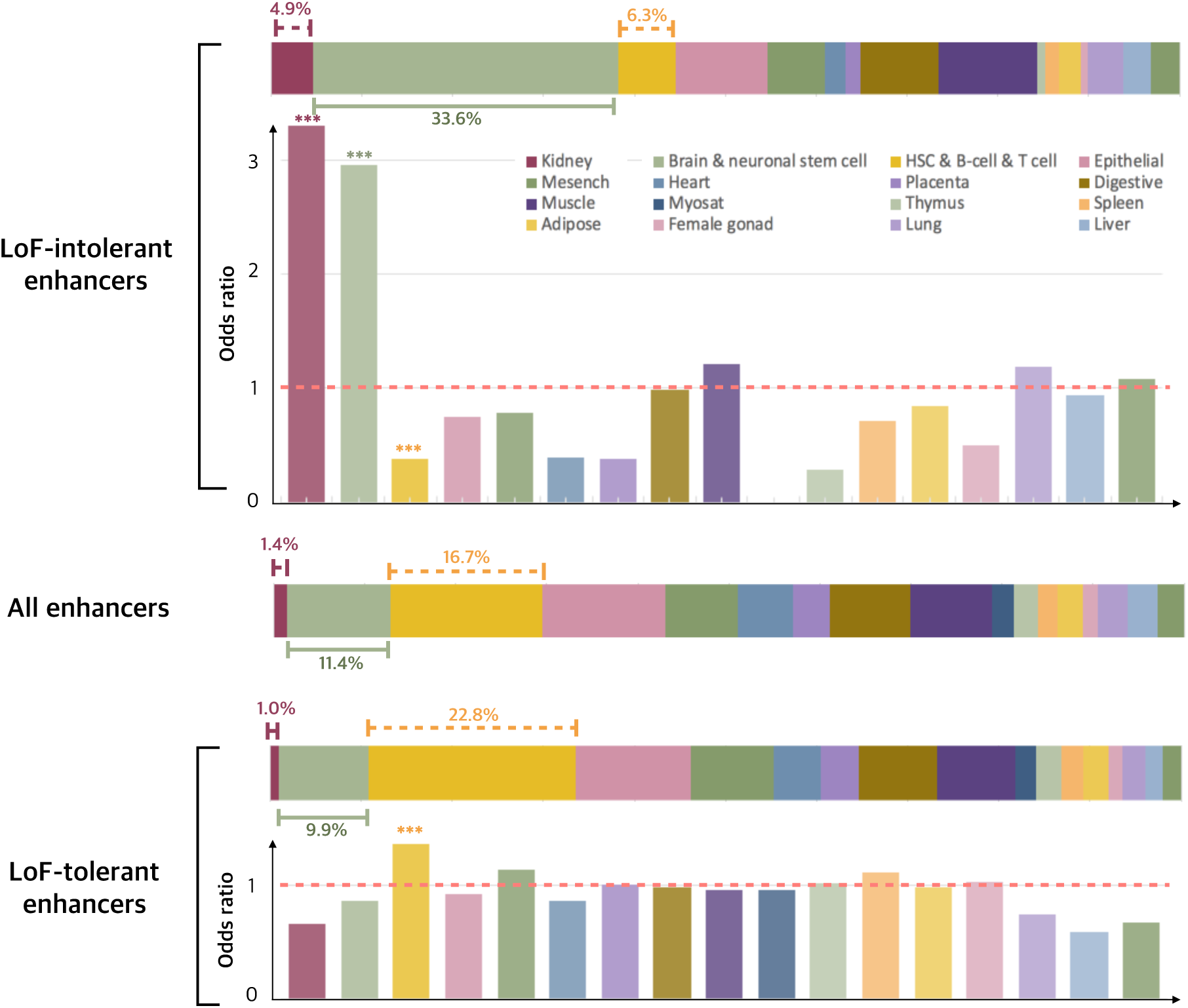
Tissue-specific enhancers. Horizontal bars show the percentage of LoF-tolerant and LoF-intolerant enhancers in each tissue type. Vertical bar plots show their odds ratios for enrichment/depletion in each tissue compared to all LoF-intolerant/-tolerant enhancers in all tissues (asterisks mark the statistical significance using Fisher’s exact test).

### Supervised learning to predict enhancer loss-of-function tolerance

Enhancer->gene regulation occurs in a complex network with interactions between enhancers and genes and among genes. Thus, to systematically predict the LoF tolerance of enhancers, we built a random forest classification model based on the network properties of enhancers and genes in the MegaNet (in total 63 features for 15 tissues as described above and in Supplementary Table 1, Methods). We also included evolutionary conservation (Siepel et al. 2005) and gene dispensability scores (Khurana et al. 2013) (Methods).

In order to avoid the prediction bias introduced by unbalanced positive and negative sample sizes, we randomly chose 50 enhancers from the LoF-tolerant enhancer set and used the 49 LoF-intolerant enhancers as the negative set to train the model. The process was repeated 50 times to sample all the 886 LoF-tolerant enhancers for training, and our model achieved an average area under the receiver operating characteristic curve (AUROC) of 0.9633 +/− 0.0002 evaluated by 10-fold cross validation on the balanced sets (Methods).

In the training process, our model uses the ultra-conserved enhancers as LoF-intolerant enhancers. Therefore, our model may bias towards enhancers residing in regions with low conservation for the predicted LoF-tolerant enhancers. In order to evaluate whether conservation alone can separate LoF-tolerant and -intolerant enhancers, we trained one model using conservation as the only feature and another model using all other features except conservation. We used one positive set with randomly selected 50 LoF-tolerant enhancers to test our models. We obtained an average AUROC of 0.92 +/− 0.0523 for the conservation-only model and 0.79 +/− 0.1801 for the other-features-only model while the final model using all features achieved an average AUROC of 0.98 +/− 0.0256 (Figure 3a). These results confirm that the integrative model gives the best performance and that the network features enable substantially better discrimination between LoF-tolerant and -intolerant enhancers than conservation alone.

**Figure 3.**
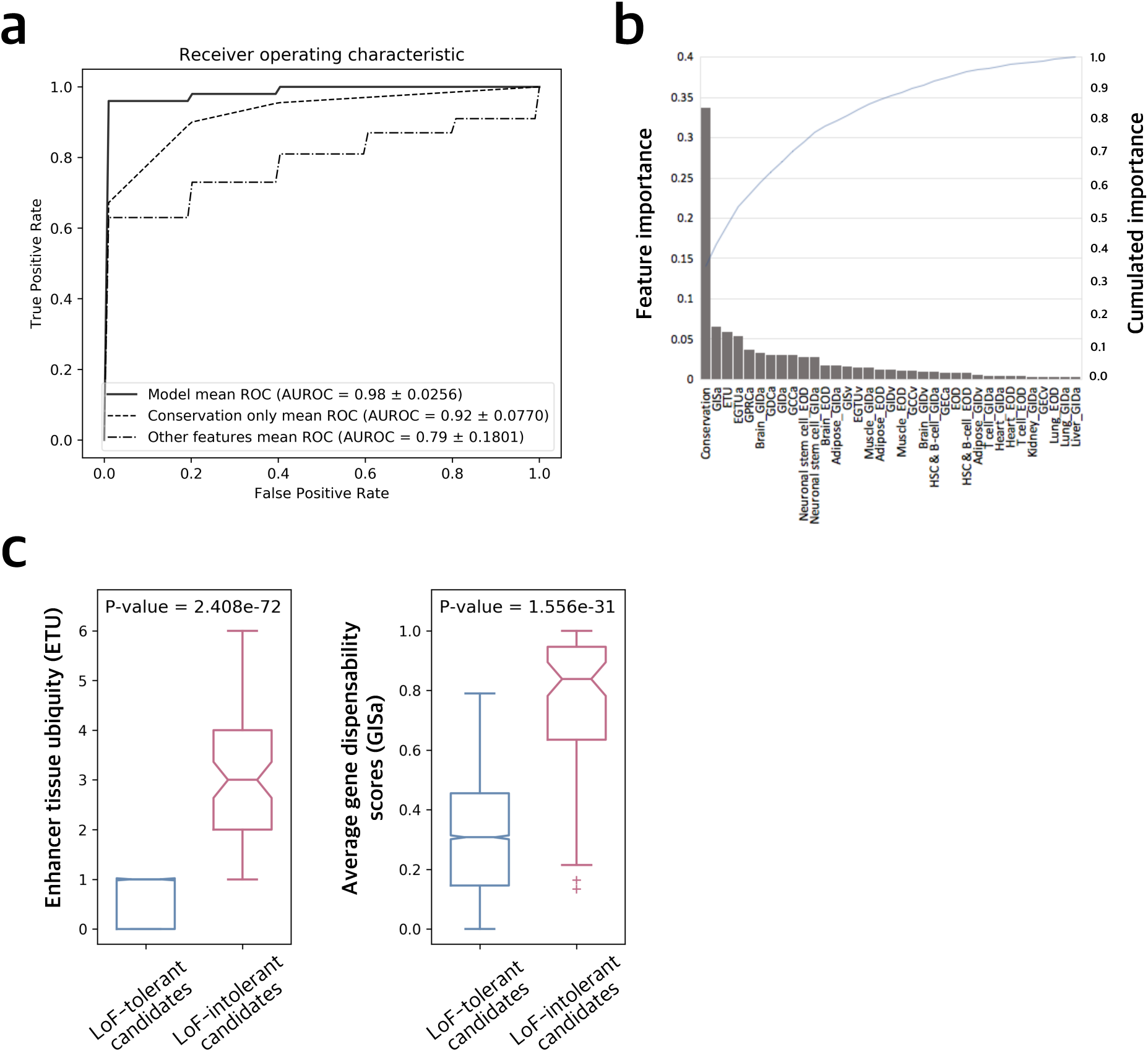
Model performance. a) 10-fold cross validation AUROC of a randomly selected set of 50 positives for the random forest classification model. b) Feature importance for the classification model. X-axis shows the features. c) Enhancer tissue ubiquity (ETU) and average gene indispensability scores (GISa) for LoF-tolerant and -intolerant enhancer candidates.

Next, we evaluated the importance of features by mean decrease impurity, which measures the decrease in the weighted impurity of the tree by each feature (Breiman 1984, Pedregosa et al. 2011). We find that while conservation is the most important feature (importance=0.3629, Figure 3b), other network properties provide substantial information to the model. On comparison of the feature importance, as expected, the features that show high importance in the model are the ones that show a significant difference between LoF-tolerant and -intolerant enhancers (as discussed above). Average edge tissue ubiquity (EGTUa) and enhancer tissue ubiquity (ETU) are the most important features after conservation, collectively contributing 18.7% of the importance. Gene indispensability scores (GIS), gene closeness centrality (GCC), out-degree of neuronal stem cell/brain enhancers and the average indegrees of genes they target also appear as important features in the model (Figure 3b).

### Prediction of novel LoF-tolerant enhancers and validation using PacBio structural variants

We applied our model on all enhancers in the MegaNet, except the ones used in training. Out of 245,093 enhancers tested, 5,677 are predicted to be highly confident tolerant to LoF with high LoF-tolerance probability (P_LoF-tol._ > 0.95), while 75 are predicted to be highly confident LoF-intolerant candidates with very low LoF-tolerance probability (P_LoF-tol._ < 0.05, Supplementary Table 3). The predicted LoF-intolerant candidates show similar patterns to the ones in the training set as they tend to be active in more tissues (P-value = 2.408e-72) and regulate genes that are more indispensable (P-value = 1.556e-31) compared to LoF-tolerant candidates (Figure 3c, Methods).

We compared the number of predicted LoF-tolerant enhancers to the predicted number of dispensable genes from Khurana et al. (Khurana et al. 2013). For comparable analysis between genes and enhancers, we took highly confident LoF-tolerant and -intolerant enhancers and used the same cut-off for gene dispensability scores (scores lower than 0.05 for dispensable and higher than 0.95 for indispensable genes). We found that the ratio of predicted LoF-tolerant to LoF-intolerant enhancers (75.7, 5677:75) is significantly higher compared to the ratio of predicted dispensable to indispensable protein-coding genes (0.483, 1259:2606, Fisher’s exact test P-value < 2.2e-16). This result is consistent with the hypothesis that LoF of genes would likely cause more severe fitness effects and must be under stronger negative selection than the LoF of regulatory elements.

Thus, in addition to the 886 homozygously deleted LoF-tolerant enhancers used in training, our model predicts additional 5,677 highly confident LoF-tolerant enhancers (P_LoF-tol._ > 0.95). We postulate that many of these enhancers have not yet been detected as LoF-tolerant because of (a) the limited sample size of whole-genome sequences and (b) undetected deletions by short-read sequencing due to the limited mappability of short reads in repetitive and complex regions. In particular, recent studies have pointed out that the map of genomic deletions with Illumina short-reads is highly incomplete. The longer sequencing reads in PacBio technology enabled the detection of many additional structural variants (SVs, including deletions), particularly in high-repeat regions (24,825 as opposed to 10,884 per human genome) (Chaisson et al. 2015, Chin et al. 2013, Kronenberg et al. 2018, Chaisson et al. 2018). We tested the performance of our method on homozygously deleted enhancers obtained from a combination of PacBio long-reads and Illumina short-reads (Chaisson et al. 2018). We found 21 novel enhancers completely deleted in a homozygous fashion in the three individuals sequenced by Chaisson et al. Our model predicted significantly higher LoF-tolerance probability scores for these enhancers than the genome average (Kolmogorov-Smirnov test P-value = 0.010, Figure 4b). This result shows that the scores predicted by our model can help with identification of LoF-tolerant enhancers even in the absence of large numbers of whole-genomes and incomplete maps of genomic deletions generated using Illumina short-reads.

**Figure 4.**
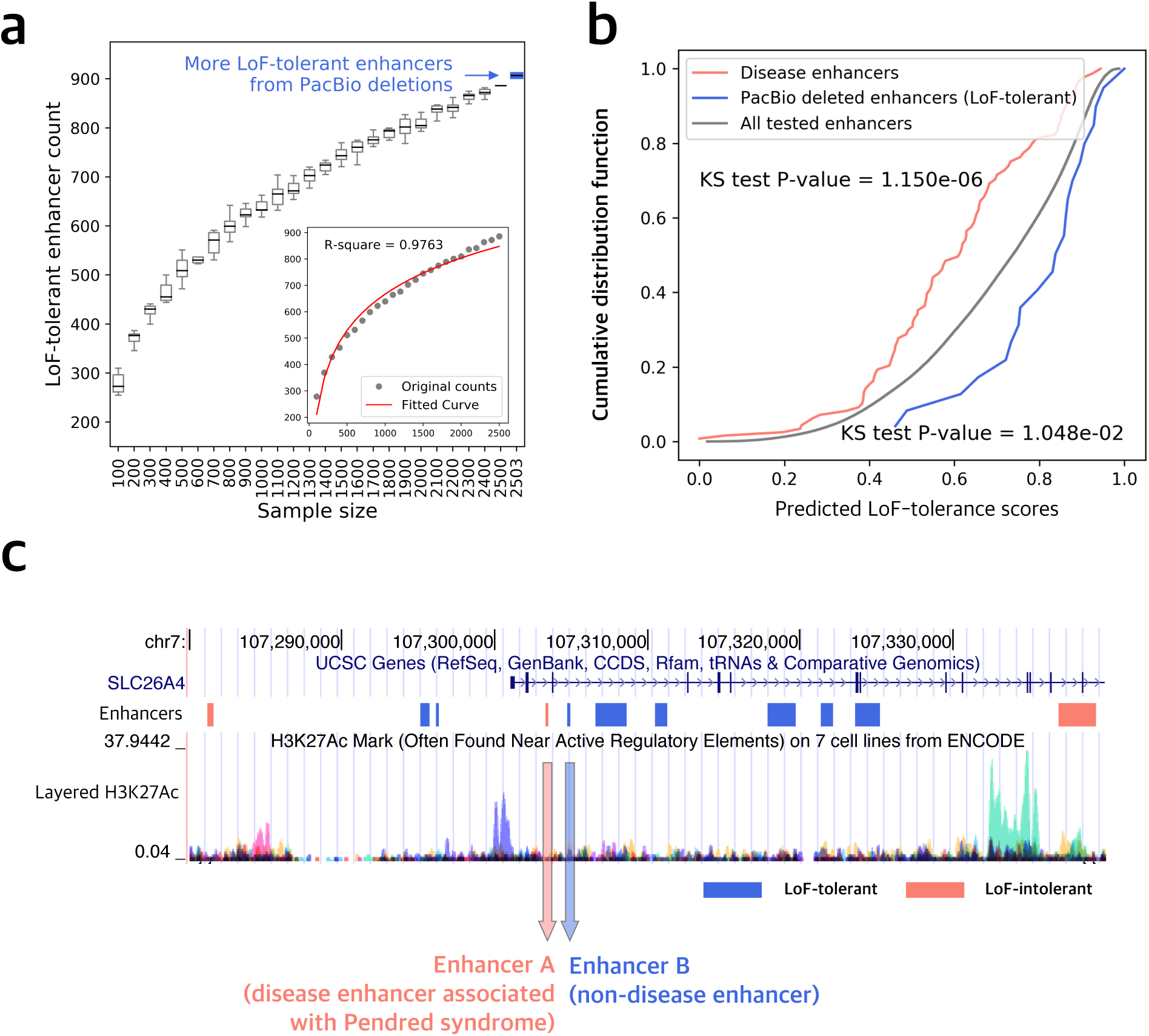
Validation using PacBio SVs and disease enhancers. a) Number of observed LoF-tolerant enhancers with increasing sample size. On the x-axis, 2503 includes the LoF-tolerant enhancers observed from 3 additional individuals sequenced using PacBio. b) Cumulative distribution function for LoF-tolerant scores for disease enhancers (red), all tested enhancers (grey), PacBio deleted enhancers (blue). KS-test P-values are between disease enhancers vs. all tested and PacBio enhancers vs. all tested. c) Genome region of *SLC26A4* and part of the enhancers regulating it. Blue denotes the predicted LoF-tolerant enhancers, while red is for predicted LoF-intolerant enhancers. Layered H3K27Ac is the modification of histone H3 lysine 27 acetylation which is associated with active enhancers.

In order to estimate how many LoF-tolerant enhancers we may expect to obtain as more whole-genomes are sequenced, we randomly chose increasing numbers of genomes in sets of 100 from 2,504 whole-genomes and calculated the number of LoF-tolerant enhancers discovered. Our power calculations using this sub-sampling approach show that the number of LoF-tolerant enhancers is likely to increase exponentially as more genomes are sequenced (Figure 4a). However, sequencing all human genomes to find all the LoF-tolerant enhancers is still infeasible even with short-reads sequencing, let alone more expensive and time-consuming long-reads sequencing. Thus, our model can serve as a practical method to predict which enhancers will be more prone to LoF-tolerance and in the interpretation of disease-associated non-coding variants as discussed below.

### Predicted LoF-intolerant enhancers and disease risk

In order to evaluate if our model can be informative for the prediction of disease-associated regulatory elements, we extracted a set of disease enhancers from DiseaseEnhancer database (Zhang et al. 2018). In this database, the authors used manual curation to collect the enhancers for which the related genetic variation has been associated with disease phenotypes or important TF binding changes (Zhang et al. 2018). We examined the LoF-tolerance scores predicted by our model for the 90 disease enhancers matched in MegaNet (Methods). We find that the disease-associated enhancers have significantly lower LoF-tolerance probabilities relative to all the enhancers (Kolmogorov-Smirnov test P-value = 1.150e-6), suggesting that our model correctly predicts their intolerance to loss of function (Figure 4b).

We further categorized these enhancers into different disease groups, for example, obesity, skin diseases, neurological disorders, artery diseases, immune disorders, and developmental diseases. We find that skin disease related enhancers have higher LoF-tolerance probability scores (Wilcoxon rank sum test P-value = 0.025, Supplementary Figure 4a), while psychological disorders related enhancers have lower LoF-tolerant scores (average predicted P_LoF-tol_ = 0.46, Wilcoxon rank sum test P-value = 0.024, Supplementary Figure 4a).

We also inspected a few prominent individual examples related to severe diseases. Previous studies have shown that a single nucleotide mutation in an enhancer regulating *SLC26A4* can cause decreased enhancer activity leading to repression of gene expression (Fuxman Bass et al. 2015), which in turn is associated with Pendred syndrome (Campbell et al. 2001, Tsukamoto et al. 2003). Pendred syndrome is a disorder associated with hearing loss caused by abnormalities of inner ear. SLC26A4 is an anion transporter and its disablement can cause hearing loss and inner ear malformation (Yang et al. 2007, Lazzereschi et al. 2005). This enhancer (Enhancer A, Figure 4c) is predicted to be LoF-intolerant by our model with P_LoF-tol._ = 0.41 (P_LoF-tol_ < 0.5), consistent with its loss of function leading to the disease. In contrast, a neighboring enhancer (Enhancer B), which is 1.2 kbp away is predicted to be LoF-tolerant (P_LoF-tol._ = 0.94). This result shows that our model can differentiate between LoF-tolerant and LoF-intolerant enhancers even when they regulate the same gene. Closer inspection of these two enhancers reveals that the reason why these two closely located enhancers are predicted to have different LoF-tolerance by our model is that Enhancer A is active in more tissues (spleen and heart) and shows higher sequence conservation (PhastCon score = 0.36) relative to Enhancer B (active in H1 embryonic stem cells with PhastCon score = 0.026, Supplementary Table 2).

In another prominent example of enhancers related to severe diseases, *ZIC3* is a protein-coding gene in the ZIC family of C2H2-type zinc finger proteins, acting as a transcriptional activator in the early stages of determining body left-right asymmetry. Mutations in *ZIC3* have been found in X-linked heterotaxy syndrome and isolated congenital heart disease (CHD) (Gebbia et al. 1997, Ware et al. 2004). Homozygous mutations in *ZIC3* in mice result in 50% embryonic lethality and live born mice exhibit severe congenital heart defects, pulmonary reversal or isomerism (Purandare et al. 2002). Out of 33 enhancers that regulate this gene, 17 are predicted to be LoF-intolerant by our model with average P_LoF-tol._ = 0.25. Previous studies have found 8 LoF mutations in coding regions of *ZIC3* related to the heterotaxy, however, they only explained ∼1% of the cases (Ware et al. 2004). Therefore, the LoF-intolerant enhancers predicted by our model may provide potential novel susceptibility loci for the study of X-linked heterotaxy and CHD.

Overall, these results suggest that the LoF-tolerance probability scores predicted by our model can provide a powerful reference for disease and clinical studies.

### Non-conserved enhancers may be LoF-intolerant

As noted above, although evolutionary conservation is the feature with the highest importance (0.338) in our model, other network features improve the performance of the model and their integration allows us to achieve an AUROC of 96%. In order to further interpret the relationship between network properties and LoF-tolerance, we examined their contribution to disease enhancers. From the disease enhancer set described in the previous section, there are 11/39 enhancers with conservation < 0.065 (median of all enhancer PhastCon scores) (Pollard et al. 2010) yet they are predicted to be LoF-intolerant by our model. One example is an enhancer regulating the gene *MFS1*. Two SNVs (rs3821943, rs4689397) in this enhancer have been associated with type 2 diabetes and shown to decrease the enhancer activity by lowering the expression of *MFS1* through luciferase reporter activity (Stitzel et al. 2010). The susceptibility loci reported in the study locate in one enhancer in our dataset with P_LoF-tol_ = 0, hence it is predicted to be a highly confident LoF-intolerant candidate. However, the conservation for this enhancer region is low (PhastCon score = 0.0043), which shows that our model can help prioritize and interpret disease variants beyond the use of evolutionary conservation (Supplementary Figure 4b).

## Discussion

In this study, we constructed a unified human regulatory network (MegaNet) by integrating tissue-specific enhancer-target networks and gene-gene interactions. To define enhancers that may be tolerant to LoF in the genome, we used deletions from 1000 Genomes Project. We describe the differences between LoF-tolerant and -intolerant enhancers in the MegaNet. We observe that LoF-tolerant enhancers regulate fewer genes and tend to be more tissue-specific. We also find that the genes regulated by LoF-tolerant enhancers tend to be regulated by more enhancers, indicating enhancer redundancy in the network. The catalogue of LoF-tolerant enhancers allowed us to develop a supervised learning method to predict the LoF-tolerance of all enhancers in the human genome using their properties in MegaNet. Independent data sets obtained using longread sequences and known sets of disease enhancers provide validation for the LoF-tolerance scores predicted by our model.

GWAS have revealed that the majority of the variants associated with complex diseases reside in non-coding regions of the genome (Hindorff et al. 2009, McCarthy and Hirschhorn 2008, Maurano et al. 2012). Moreover, whole exome sequencing could only find the causal variants for ∼25-50% of patients (Wortmann et al. 2015, Kremer et al. 2017). It is likely that regions excluded from exome sequencing, namely non-coding regions, harbor the variants explaining many of the remaining unexplained cases (Valente and Bhatia 2018). Major international efforts such as the UK Biobank and TOPMed (NHLBI Trans-Omics for Precision Medicine) aim to use whole-genome sequencing to uncover disease variants (Bycroft et al. 2018, Turnbull et al. 2018, Sarnowski et al. 2018, He et al. 2019, Telenti et al. 2016, Perkins et al. 2018). The LoF-tolerance scores for enhancers provided here can significantly facilitate the interpretation and prioritization of non-coding sequence variants in whole-genome sequencing studies.

## Materials and Methods

### Obtaining enhancer-gene networks

Enhancer-gene networks in different tissues were obtained from the ENCODE+Roadmap LASSO dataset in Cao et al. (Cao et al. 2017, http://yiplab.cse.cuhk.edu.hk/jeme/). In Cao et al, they collected ChIP-seq data for H3k4me1, H3K27ac, H3K27me3, DNase-seq together with ChromHMM-predicted active enhancers to generate a union set of enhancers. In total, we collected 246,028 unique enhancers regulating 19,170 genes from enhancer-target networks from all tissue types. We grouped 127 Roadmap tissue types by the given sample group into 19 tissue groups and discarded ungrouped cell types.

In order to identify LoF-tolerant enhancers, we first identified all deletions existing in a homozygous state in any one individual in the 1000 Genomes Phase 3 data (Sudmant et al. 2015). We excluded any deletion overlapping coding exon regions and then intersected the remaining deletions with enhancer coordinates to obtain our list of 886 LoF-tolerant enhancers. Only enhancers that are 100% deleted were included.

In order to identify LoF-intolerant enhancers, we started with ultra-conserved elements and retained only those showing consistent reporter gene expression (Bejerano et al. 2004, Visel et al. 2007, Visel et al. 2008, Dickel et al. 2018). We intersected the remaining elements with enhancer coordinates in our dataset, keeping only those with >50% reciprocal overlap. In total, we define 49 LoF-intolerant enhancers.

We compared the length distributions of enhancers and deletions (Supplementary Figure 5). The average length of deletions is much longer than enhancers. Thus, LoF-tolerant enhancers are likely not biased towards shorter enhancers (shorter enhancers are more likely to be completely deleted). To be more stringent, we still excluded the length of enhancers as a feature in the following analysis.

### Tissue-specific subnetworks

To distinguish enhancer activity differences between tissues, we extracted tissue-specific networks from the MegaNet. Enhancers in HSC & B-cell and Epithelial tissues exhibit significant differences in tissue-specific network properties between LoF-tolerant and LoF-intolerant enhancers (Wilcoxon rank sum test P-value < 0.05, Supplementary Figure 3b).

### Collecting features for the model

Besides the tissue specificity information of enhancers, we also used the gene centralities and gene indispensability scores (Khurana et al. 2013) as measurements for gene priority in the network. In order to only consider the direct interactions between gene pairs, indirect interactions, genetic interaction and regulatory interactions, were excluded from our integrated network. Enhancer-target network features were calculated using Python networkX package (Hagberg, Swart and S Chult 2008). Conservation scores for sequence were obtained from PhastCons (Pollard et al. 2010).

Detailed information about network features is provided in Supplementary Table 1. The enhancer tissue ubiquity (ETU) is the total number of tissues that the enhancer is active in. The enhancer-gene edge tissue Ubiquity (EGTU) is the number of tissues that the enhancer-gene regulation edge is active in. For enhancers that regulate multiple genes, to transform gene features for those regulated genes into an enhancer feature, we took both the average and variance for each gene features and represented it with extension “a” (average) or “v” (variance). For each enhancer, we denote ETU as n, then EGTU is a list of (*e*_1_, *e*_2_, …, *e*_*n*_). The EGTUa will be 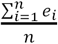,and the EGTUv is 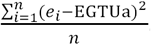.

### Feature selection

To avoid overfitting introduced by features correlated with each other, we calculated the Spearman distance between each feature. We noticed that features for tissue type adipose/epithelial and digestive are strongly correlated with each other, thus only one of them (adipose) was kept for further model building. In addition, tissue type myosat and mesench are mixed with other tissue clusters, so we eliminated them from the final tissue set. In the end, there are in total 15 tissue types considered and 62 features overall.

### Model building and testing

The model was built using tools from Python Scikit-learn package (Pedregosa et al. 2011). Random and grid searches were performed to find the best parameters for the random forest classifier. 10-fold cross validation was performed to evaluate the performance of the model. We selected 50 LoF-tolerant and 49 LoF-intolerant enhancers to train the classifier. To avoid overfitting, we repeatedly sampled through all LoF-tolerant enhancers 50 times. The mean AUROC is 0.9633 +/− 0.0002. Due to the small sample size of LoF-intolerant enhancers, we also randomly chose 50 enhancers from neither the LoF-tolerant nor intolerant set as “LoF-intolerant” to test overfitting of the model. We performed the same parameter searching and cross validation repeatedly 50 times and obtained mean AUROC of 0.6154 +/− 0.0697, indicating that the small sample size for LoF-intolerant enhancers did not lead to overfitting.

We applied the model on all other enhancers in the network and predicted their probability to be LoF-tolerant as their LoF-tolerant scores. The predicted LoF-tolerant probabilities are the mean predicted class probabilities of the trees in the forest (Pedregosa et al. 2011). Among 245,361 enhancers tested, 194,812 (P_LoF-tol_ >= 0.5) are predicted to be LoF-tolerant enhancers, while 50,281 to be LoF-intolerant (P_LoF-tol_ < 0.5) and 5,677 are predicted to be highly confident LoF-tolerant (P_LoF-tol._ > 0.95), while 75 are predicted to be highly confident LoF-intolerant candidates with very low LoF-tolerance probability (P_LoF-tol._ < 0.05).

### Validation

To further validate our observation that there are additional LoF-tolerant enhancers in human genomes, we obtained novel deletions to identify LoF-tolerant enhancers. Those novel deletions were from the 1000 Genomes structural variation consortium where they used integrated structural variation calling methods including both Illumina short reads and PacBio long reads sequencing for three individuals from 1000 Genomes trio studies (Chaisson et al. 2018). In total, we used 12,939 deletions from the PacBio structural variants set that were present in the three children from the trio family and intersected them with 1000 Genomes Phase 3 deletions. There are 11,118 novel deletions with less than 80% overlap with the 1000 Genomes Phase 3 deletions. Out of those novel deletions, 21 of them can delete enhancers completely from our enhancer set.

### Disease enhancers

Disease enhancers were collected from Zhang et al. (Zhang et al. 2018). We intersected our enhancers with the 1,059 disease enhancers which defined in Zhang et al., if no overlap found then take the closest neighbored enhancer. After this, keep only the disease enhancers that its target gene from the DiseaseEnhancer matches the enhancer-gene regulation from our dataset. To further filter out the disease enhancers related to somatic variants, we excluded enhancers associated with cancer. In the end, we collected 90 enhancers in our dataset with disease associations.

## Figures and Supplementary Tables

**Supplementary Figure 1.**
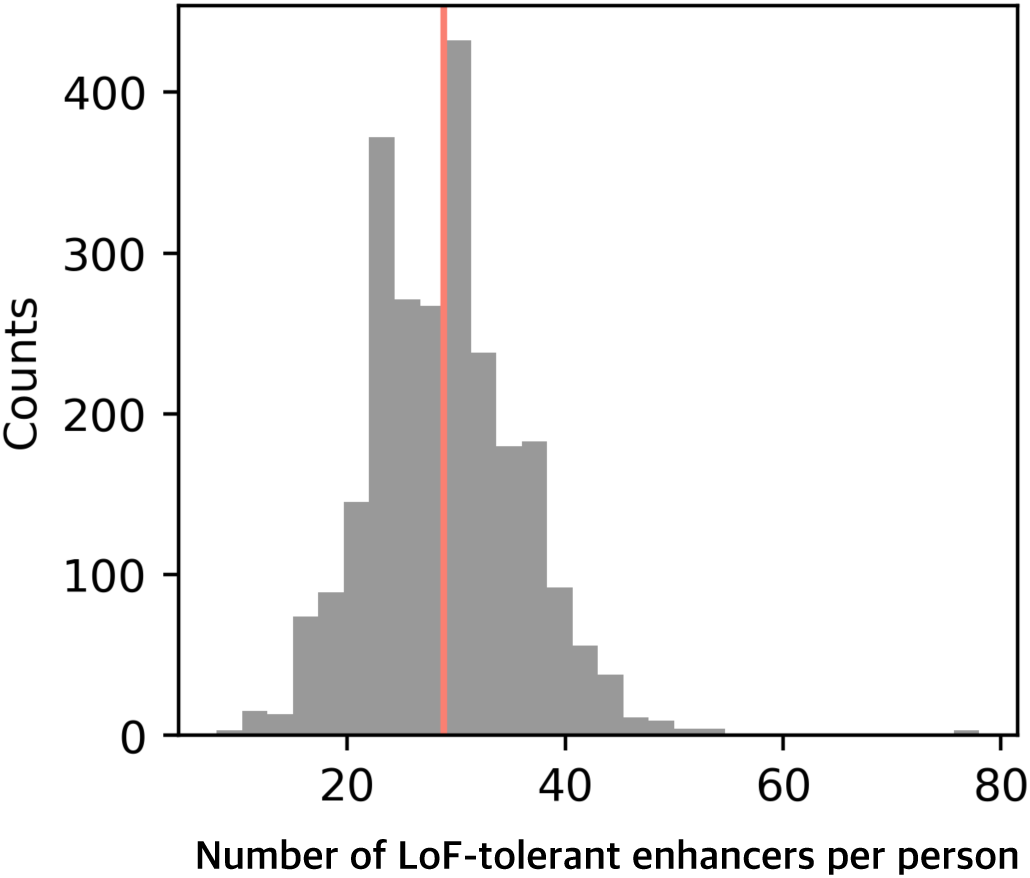
Number of LoF-tolerant enhancers per individual from 2,504 genomes. Each individual has on average 28 enhancers (red vertical line) completely and homozygously deleted in the genome.

**Supplementary Figure 2.**
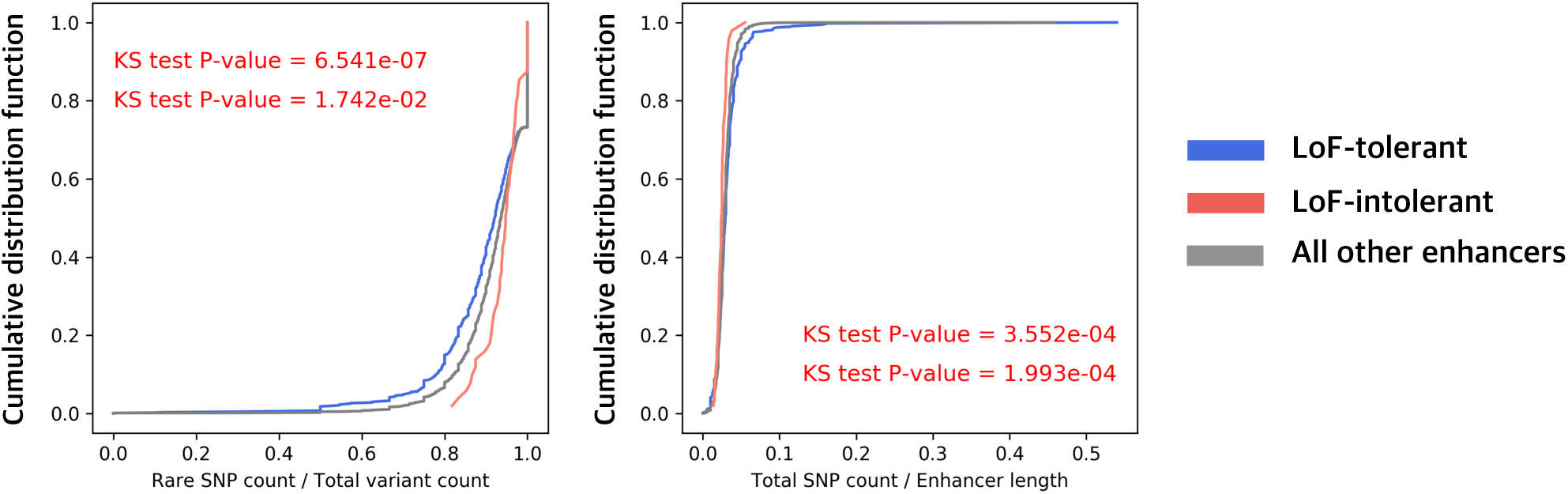
Comparison of enrichment of rare variants and all polymorphisms between LoF-tolerant and -intolerant enhancers and all other enhancers (genome-wide, GW). Upper P-value is for LoF-tolerant vs. GW, while lower P-value is for LoF-intolerant vs. GW. The P-values were calculated by Kolmogorov-Smirnov test (KS test).

**Supplementary Figure 3.**
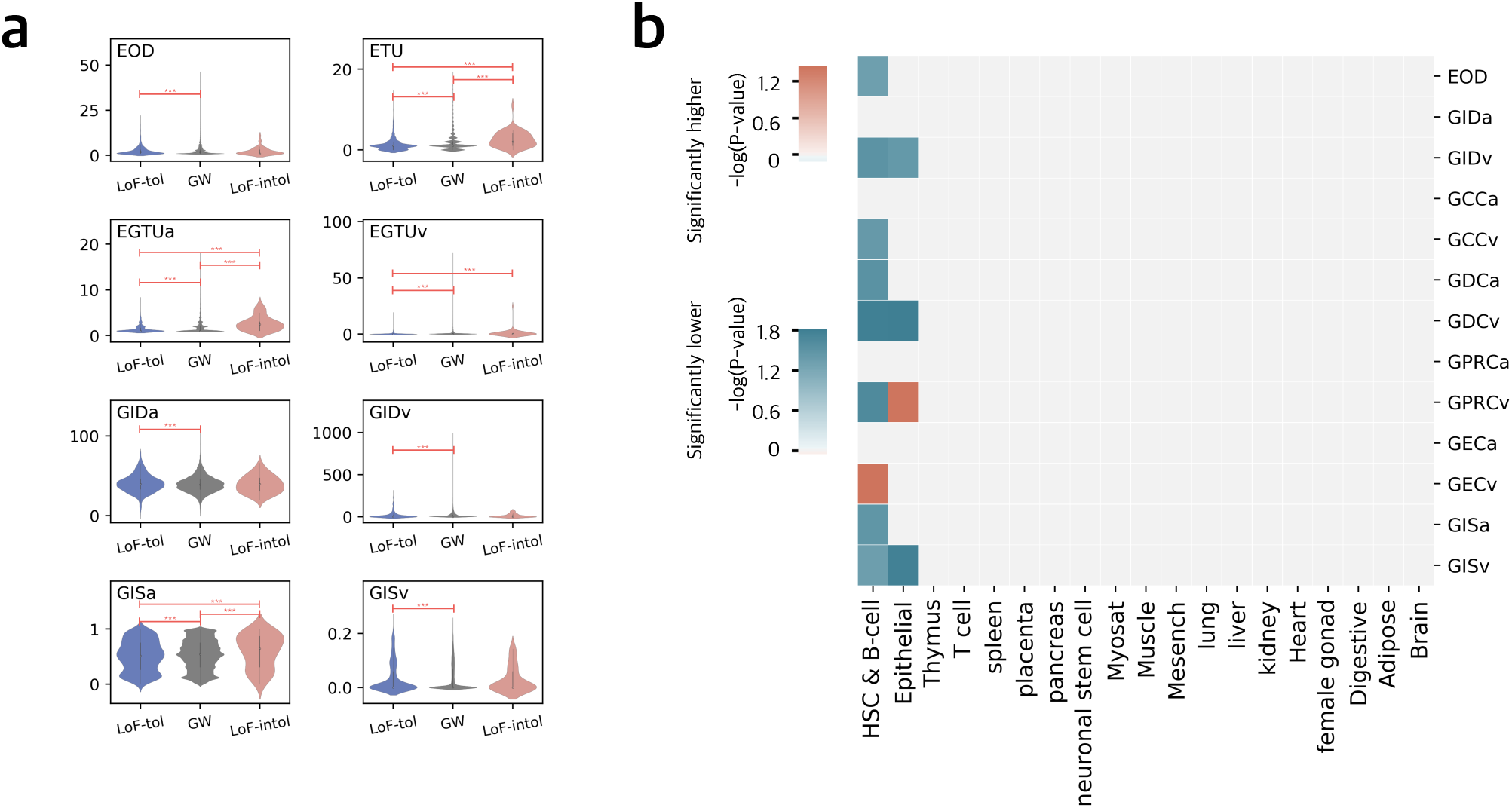
Network features in the MegaNet and in tissues-specific networks. a) network features in the MegaNet, significant comparisons are marked by asterisks. b) Each column represents a tissue-specific network comparison between LoF-tolerant vs. LoF-intolerant enhancers (P-values were computed using Wilcoxon rank sum test).

**Supplementary Figure 4.**
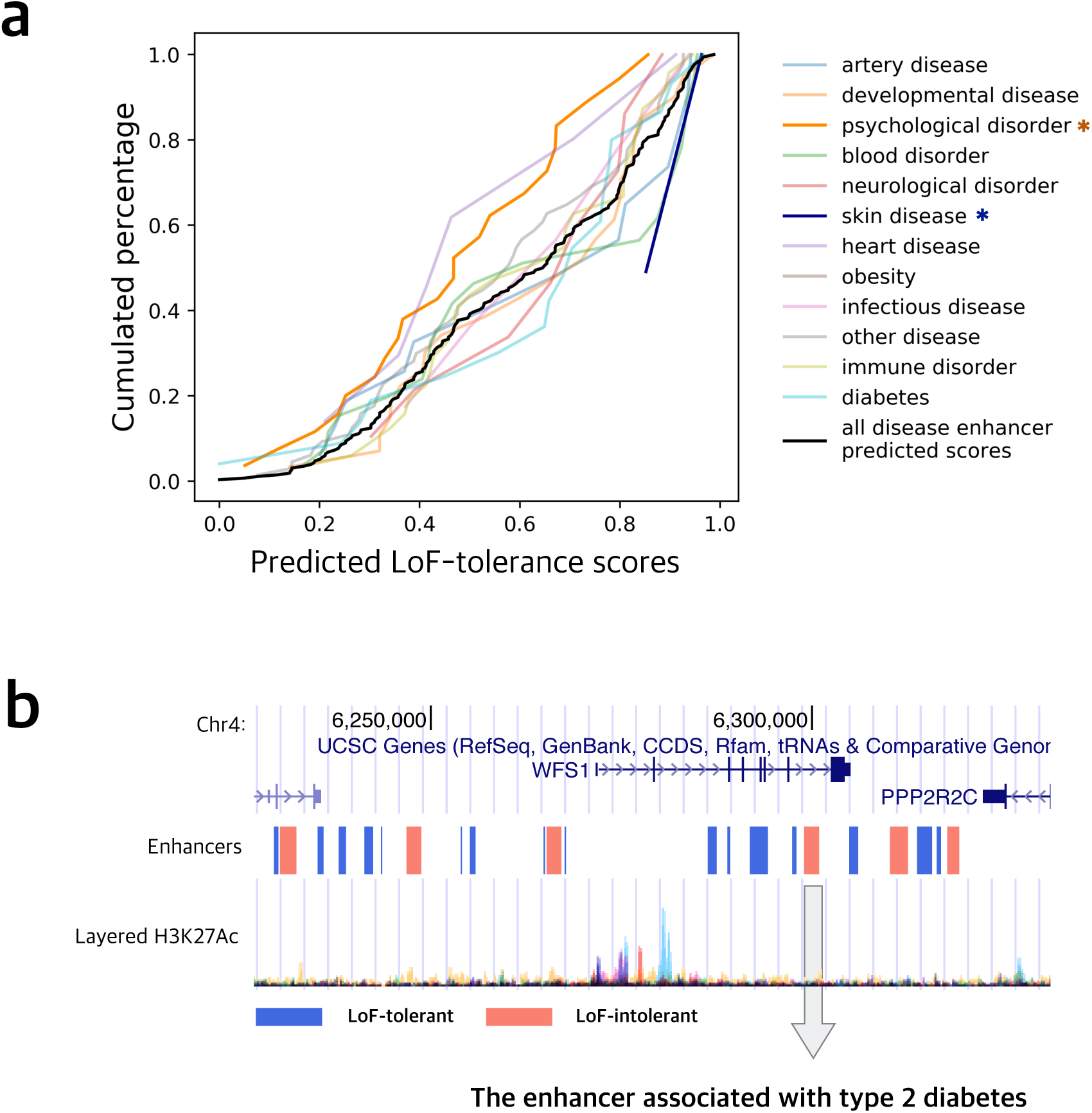
Disease enhancers. a) Predicted LoF-tolerance scores for disease enhancers by disease types. Y-axis is the cumulated percentage of enhancers for the corresponding LoF-tolerance scores on x-axis. Disease types are colored as shown, significant ones (P-value < 0.05) are marked by asterisks. b) Genome region of *WFS1* and part of the enhancers regulating it. Blue denotes the predicted LoF-tolerant enhancers, while red is for predicted LoF-intolerant enhancers. Layered H3K27Ac is the modification of histone H3 lysine 27 acetylation which is associated with active enhancers.

**Supplementary Figure 5.**
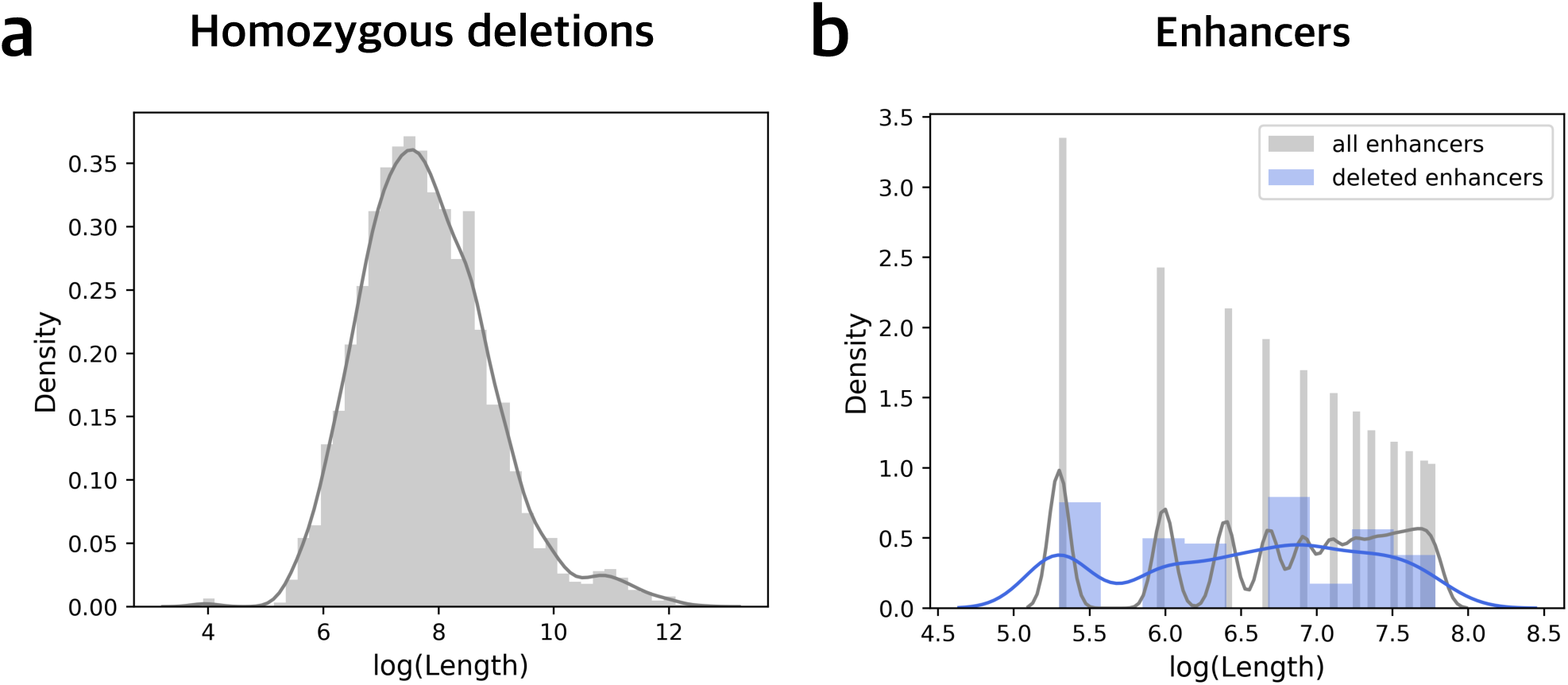
Length distribution of homozygous deletions, deleted enhancers and all enhancers.

**Supplementary Table 1.** Summary of network features.

**Supplementary Table 2.** Categories of ENDODE and Roadmap tissues

**Supplementary Table 3.** Predicted LoF-tolerance scores for all enhancers in this study.

